# The impact of tail-cup handling as an alternative method for picking up laboratory mice

**DOI:** 10.64898/2026.11.16.766753

**Authors:** Ryan Nolan, Matthew Leach, Candy Rowe, Jane L. Hurst

## Abstract

The way that laboratory rodents are handled has a significant impact on their welfare. Picking up mice by the tail increases anxiety, reduces voluntary interaction with handlers, and reduces behavioural test reliability compared to more refined handling methods, such as picking up mice inside a handling tunnel or cupping freely on the hand. An alternative in use by the community is to restrain a mouse by the tail-base while scooping it up on the hand and supporting its weight (tail-cup method). Here, we assess whether tail-cup handling is a welfare refinement compared to picking up mice by tail or tunnel, using both sexes of two strains (BALB/c, C57BL/6). Tail-cup handling reduced aversion to interact with the handler compared to tail handling, but to a much smaller extent than tunnel handling. Tail-cup and tunnel handling both reduced anxiety in an elevated plus maze relative to tail-handled mice. However, test reliability was substantially better using tunnel handling in a habituation-dishabituation task, greatly reducing numbers of animals required for testing. These findings generalised across sexes and strains tested. While tail-cup handling provides some improvement over traditional tail handling, tunnel handling appears to be more beneficial for mouse welfare and behavioural reliability.

## Introduction

Where no suitable replacements are available, laboratory animals continue to be used for biomedical and biological research; however, their welfare must be a primary consideration. This is underpinned by legislation governing the use of animals in research both nationally and internationally, which requires that no unnecessary harm is caused to animals. In addition to eliminating negative experiences as far as practicable, the benefits of positive experiences among animals are increasingly recognised ^1,2^. Housing and husbandry procedures, together with interactions with carers and researchers, are likely to comprise the majority of laboratory animals’ lifetime experiences. Animals need to be handled for a wide range of procedures and basic husbandry, so it is essential that this experience involves as little stress and fear as possible and, where possible, establishes a positive relationship between animal and handler.

Research has revealed that the methods used to pick up laboratory rodents, which are essential for handling or transferring animals outside their cages, have a substantial impact on indicators of negative or positive animal welfare ^3^. Traditionally, the standard method to pick up laboratory mice has been to grasp and lift them by the base of the tail ^4,5^, termed tail handling. Once the animal has been lifted from the cage, its weight is usually supported on a surface (such as the hand, arm, or cage lid) while still restrained by the tail, although an animal’s weight may not be supported for very brief lifts, such as the transfer of animals between cages ^5^. However, there is now substantial evidence that tail handling has a negative impact on animals when compared to other more recently developed methods, such as the use of a handling tunnel (usually a plastic tube present in the home cage ^6^), home cage ladders ^7^, or cupping animals on the palm of the hand without restraint ^3,8,9^. For example, mice picked up by the tail show greater anxiety-like behaviours in elevated plus maze and open field tests than those accustomed to handling by tunnel, hand cupping or cage ladders ^3,6–12^. Experience of tail handling in mice induces aversion towards the handler, with animals consistently avoiding interaction with handlers even after multiple days or weeks of familiarisation ^9,13,14^. By contrast, mice picked up by handling tunnel or ladder interact readily with their handling device and their handler, seeming to seek the opportunity for handler interaction once they are familiarised with the experience of being lifted. It should be noted, though, that two studies used a much more prolonged tunnel handling technique that did not direct mice straight into a tunnel. In both cases, the time taken to handle mice using a tunnel was substantially increased, and these studies failed to see a consistent difference between tunnel and tail handling ^15,16^. Other studies have shown that tail handling also increases depression-like behaviour ^10,14,17^, reduces resilience against negative events ^13^, and can increase litter mortality in breeding stock that are tail-lifted by forceps compared to animals picked up using a handling tunnel ^18^. Thus, picking up mice by the tail appears to have a broad range of negative impacts on animal welfare compared to alternative refined methods. Further, it has long been recognised that background stressors in the animal’s environment can negatively impact the quality of animal-based research, for example by increasing variability or inducing abnormal responses, which can lead to inappropriate conclusions being drawn ^19,20^. In accordance with this, studies have shown that picking up mice by the tail significantly reduces the reliability of behavioural tests that are sensitive to the effects of stress, such as willingness to explore and investigate test stimuli ^21,22^ or increased variation in measures of response to pharmacological manipulation ^21,22^.

Currently, it is not clear why picking up mice by the tail induces such negative responses, which are not apparent if animals are just restrained by the tail after being picked up by refined handling methods ^3^. Being lifted and suspended solely by the tail while unsupported might be uncomfortable or painful, even if only for a brief moment. However, as yet, there is no evidence that picking up healthy mice by the tail is painful based on grimace scoring ^23^. More prolonged suspension by the tail is clearly stressful and has been used as an inescapable stress stimulus in research on depression ^24^ and to trigger epilepsy in susceptible animals ^25^. It is also possible that animals respond poorly to being approached from behind when captured and restrained by the tail, potentially stimulating an ancestral imperative to avoid capture and flee from predators ^3^; their natural response to pull away is prevented, and they are lifted up backwards in a relatively helpless and vulnerable position. This contrasts with tunnel, hand cupping, ladder handling, or use of an upturned shelter ^26^, where the animal is approached from the front (or side) and is allowed to step into or onto the handling device. This active engagement of the animal with the handling process may induce more positive responses, even though alternative escape routes are blocked and mice are guided efficiently to the handling device ^27^.

An alternative method of handling currently used in the community combines tail handling with cupping on the hand, as demonstrated in responses to a survey by Henderson and colleagues ^28^. After grasping the base of the tail between thumb and forefinger, the mouse is scooped up on the same hand so that it is lifted on the hand but restrained by the tail, a method that we refer to here as the tail-cup method. As far as we are aware, the impacts of this method on mouse welfare have not yet been formally assessed. This method might avoid the negative responses associated with traditional tail-handling methods if these are caused by tail suspension, as the animal’s weight is fully supported before lifting. Evidence shows that animals scooped onto the hand without restraint (cupping) generally show much more positive responses than those picked up by tail ^3,8,21^. However, restraining by the tail in the tail-cup method has a practical advantage, as animals cannot jump from the hand if they choose to. Nonetheless, tail-cup handling still involves approach and capture from behind, which may limit the benefits if a more cooperative approach to handling is key to the animal’s welfare.

In this study, we compare the responses of mice to repeated but very brief pick-up by tail, tail-cup, or tunnel methods, using both sexes of two common mouse strains (BALB/c and C57BL/6, shortened to B6) with contrasting behavioural phenotypes ^29–33^. BALB/c mice are generally reported to be more stress vulnerable than B6 mice, displaying higher risk assessment and avoidance while B6 are more explorative and risk taking. After experience with their assigned handling method, we assessed willingness to approach a handler in the home cage and anxiety in an elevated plus maze test. We also evaluated their performance in a habituation-dishabituation test that assessed their recognition of a novel conspecific urine scent, requiring normal behaviour to explore, investigate and remember scent from a different strain.

## Results

### Voluntary Interaction after Handling

To assess willingness to interact with a handler after experience of handling by their assigned method, we conducted two types of voluntary interaction test. First, we assessed response to the handling device itself (gloved hand holding a tunnel for the tunnel group, or gloved hand only for tail and tail-cup groups) held unmoving in the cage for 60s after the first and ninth handling sessions. To compare response to handler contact, we then also assessed response of all three treatment groups to the gloved hand only (see Methods for further details and study timeline).

From the very first handling session, handling method strongly influenced willingness to interact with the handling device (F_2,34_ = 43.00, P < 0.0001; **Table 1**). Mice handled using a tunnel spent substantially more time interacting with their handling device than those handled by tail (P < 0.0001) or by tail-cup (P < 0.0001), while there was no difference between tail and tail-cup (P = 0.56; **Table 1**; **Fig. 1a**). Neither sex nor strain significantly influenced these responses.

**Table 1.** Effects of handling method, sex, and strain on willingness to interact voluntarily with the handler (GLM ANOVAs, data summarised in **Fig. 1**).

| Voluntary Interaction | Handling Session 1 |  | Handling Session 9 |  | Gloved Hand |  |
|---|---|---|---|---|---|---|
| Handling | F<sub>2,34</sub> = 43.00 | **P < 0.0001** | F<sub>2,34</sub> = 99.03 | **P < 0.0001** | F<sub>2,37</sub> = 40.79 | **P < 0.0001** |
| Sex | F<sub>1,34</sub> = 0.15 | P = 0.71 | F<sub>1,34</sub> = 0.83 | P = 0.37 | F<sub>1,37</sub> = 0.44 | P = 0.51 |
| Strain | F<sub>1,34</sub> = 1.16 | P = 0.29 | F<sub>1,34</sub> = 3.30 | P = 0.08 | F<sub>1,37</sub> = 1.44 | P = 0.24 |
| Handling x Sex | F<sub>2,34</sub> = 3.21 | P = 0.053 |  |  |  |  |
| Post hoc comparisons: |  |  |  |  |  |  |
| Tail vs Tail-Cup |  | P = 0.56 |  | **P = 0.005** |  | **P = 0.007** |
| Tail vs Tunnel |  | **P < 0.0001** |  | **P < 0.0001** |  | **P < 0.0001** |
| Tunnel vs Tail-cup |  | **P < 0.0001** |  | **P < 0.0001** |  | **P < 0.0001** |

**Figure 1.**
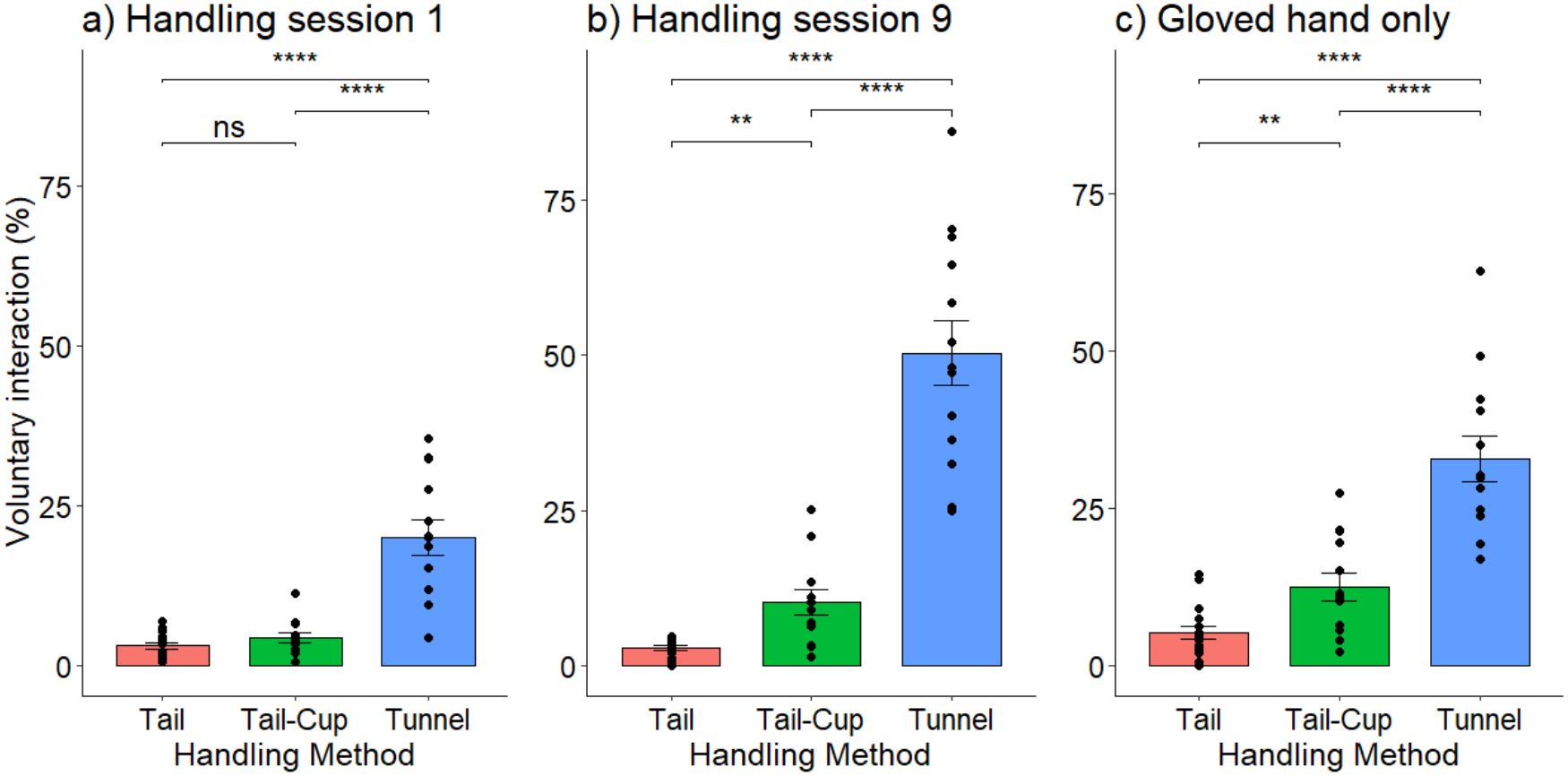
Voluntary interaction among mice handled by one of three different methods (mean proportion of 60s test time averaged across mice in the same cage ± SEM, data pooled across sexes and strains). Time spent interacting with handling device (gloved hand for tail and tail-cup groups, gloved hand holding tunnel for tunnel group) following (**a**) first handling session and (**b**) ninth handling session. (**c**) Time spent interacting with gloved hand only following an additional handling session. P values from Tukey post hoc tests (ns P > 0.05, **P < 0.01, ****P < 0.0001), see **Table 1** and **Supplementary Table 1** for full ANOVA analyses.

By the ninth handling session, handling method continued to have a substantial effect on interaction with the handling device (F_2,34_ = 99.03, P < 0.0001; **Table 1**). The difference between methods was now even stronger than in the first session (interaction between handling method x session: F_2,35_ = 20.14, P < 0.0001; **Fig. 1a,b**). Mice handled by tunnel (P < 0.0001) or by tail-cup (P = 0.005) were both more willing to interact than tail-handled mice, but interaction was much more prolonged among those handled by tunnel than by tail-cup (P < 0.0001; **Fig. 1b**). Repeated experience of tunnel handling promoted prolonged voluntary interaction (session 9 vs 1: F_1,12_ = 39.60, P < 0.0001), with these mice now spending half the test time on average interacting with the tunnel and hand. Interaction increased a little with experience among tail-cup mice (session 9 vs 1: F_1,11_ = 6.67, P = 0.025), but there was no increase among tail-handled mice (F_1,13_ = 0.001, P = 0.97; **Fig. 1b**). Neither sex nor strain significantly influenced these responses (**Table 1**).

Once familiar with handling, we also compared voluntary interaction with a gloved hand alone across all three handling groups. Again, handling method had a strong influence on willingness to interact (F_2,37_ = 40.79, P < 0.0001) while sex or strain had no effect (**Table 1**). Interaction with a gloved hand was much more prolonged for mice handled by tunnel than tail-cup (P < 0.0001) or tail (P < 0.0001), while tail-cup-handled mice continued to interact more than those picked up by the tail (P = 0.007; **Fig. 1c**).

### Anxiety in an Elevated Plus Maze Test

The effects of the different handling methods on anxiety were assessed using an elevated plus maze (EPM) test. More anxious animals spend less time on the two open arms of the four-armed maze, make fewer entries to the open arms and show more protected stretch attend postures (PSA) where the front half of the body stretches out onto an open arm, but the animal then withdraws without entry ^34,35^. Handling method influenced both time spent on the open arms of the maze (F_2,35.1_ = 7.53, P = 0.002) and number of open arm entries (F_2,32.8_ = 4.83, P = 0.015; **Table 2)**. Tail-handled mice were the most anxious, spending less time on the open arms than both tunnel- (p = 0.003) and tail-cup-handled mice (p = 0.012; **Fig. 2a; Table 2**), and entering the open arms significantly less frequently than those handled by tunnel (p = 0.027) or by tail-cup (p = 0.035; **Fig. 2b; Table 2**). Neither measure differed between tunnel- and tail-cup-handled mice. Both sexes and strains showed similar responses to handling method (no significant interactions), although, overall, BALB/c mice spent longer on open arms than B6 mice (162.2 ± 7.9 vs 117.0 ± 7.0s, P < 0.0001), while B6 mice entered open arms more frequently than BALB/c (17.3 ± 1.3 vs 12.9 ± 0.7, P = 0.03).

**Table 2.**
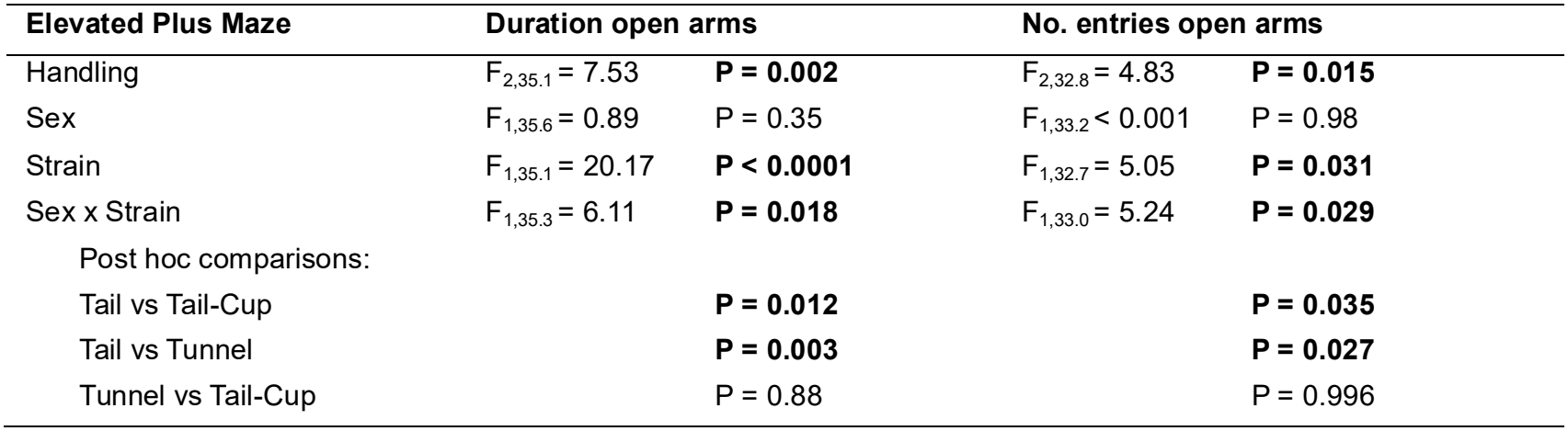
Effects of handling method, sex, and strain on anxiety in the elevated plus maze test (GLMM ANOVAs, data summarised in Fig. 2).

**Figure 2.**
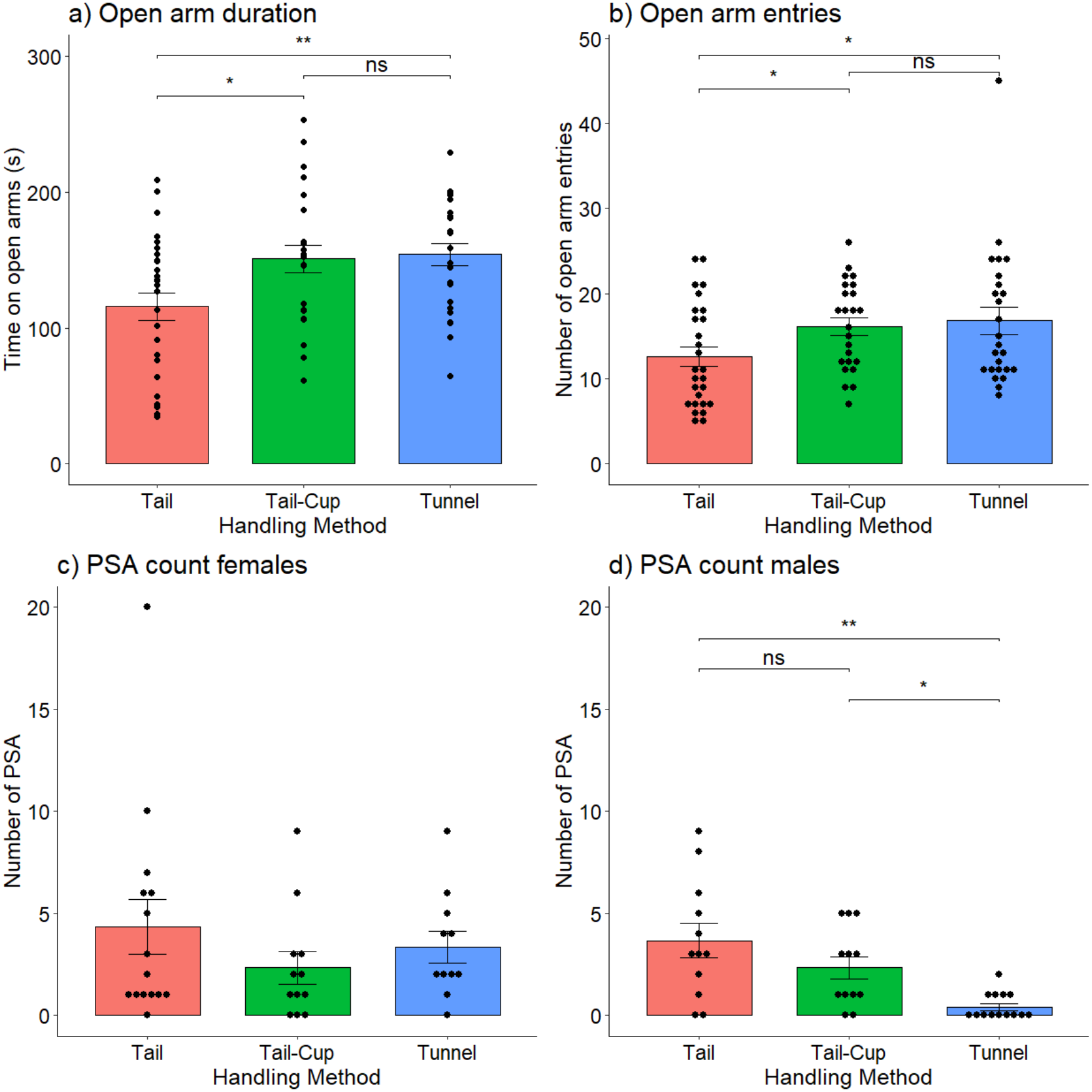
The effects of handling method on anxiety in the elevated plus maze (mean ± SEM, data pooled across sexes and strains unless otherwise stated). **a**) Duration of time on open arms. **b**) Number of open arm entries. The number of protected stretch attend (PSA) postures by (**c**) females and (**d**) males. P values from Tukey post hoc tests (ns P > 0.05, *P < 0.05, **P < 0.01), see **Table 2** and **Supplementary Tables 2&3** for full ANOVA analyses.

The effect of handling method on the number of protected stretch attend postures (PSA count) on the EPM differed between the sexes (handling x strain: F_2,33.0_ = 3.41, P = 0.045; **Supplementary Table 3**), so separate models were fitted for each sex. PSA count was not influenced by handling method in females (F_2,16.4_ = 0.95, P = 0.41; **Figure 2c**) but differed among males (F_2,14.4_ = 8.11, P = 0.004; **Figure 2d**). Both tail (P = 0.003) and tail-cup-handled males (P = 0.039) exhibited a greater PSA count than tunnel-handled males, with no difference between tail-cup and tail-handled males. PSA count was not influenced by strain (**Supplementary Table 3**).

EPM control measures (time spent in closed arms and closed arms entries) were not influenced by handling method or sex. These measures differed between strains (analysis in **Supplementary Table 4**).

### Behavioural Reliability in a Habituation-Dishabituation Test

We assessed the effect of handling method on reliability in a behavioural test that required mice to investigate and recognise a difference between two distinct conspecific urine stimuli in a habituation-dishabituation paradigm. Mice were placed into an arena with a streak of unfamiliar male urine (from a different strain to their own) and allowed to investigate. This was repeated for three five-minute trials, allowing mice to become familiar with the scent. On the fourth trial, the male urine stimulus was from another strain. If mice recognise that the two scents differ, they should investigate the novel scent in the fourth trial for longer than the familiar scent in the third trial (known as the dishabituation response) ^21^.

Handling method had a highly significant effect on the dishabituation response (Kruskal-Wallis test, H = 21.45, 2df, P < 0.0001; **Figure 3**). This was substantially stronger and more reliable in mice that had been handled by tunnel, compared to those handled by tail (P < 0.0001) or by tail-cup (P = 0.003). By contrast, the response did not differ between tail or tail-cup methods. These responses were consistent across sexes and across strains.

**Figure 3.**
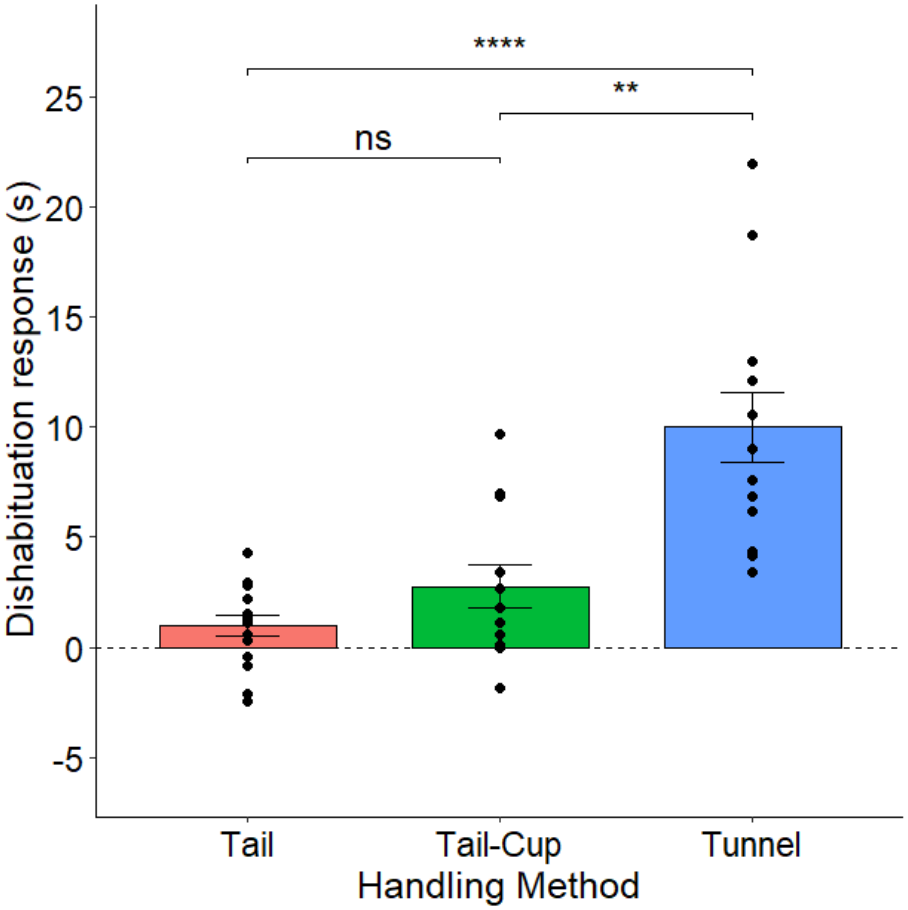
Effects of handling methods on dishabituation response (trial 4 – trial 3 stimulus investigation, mean ± SEM, data pooled across sexes and strains). P values from post hoc Dunn tests (ns P > 0.05, **P < 0.01, ****P < 0.0001).

We also examined whether mice achieved a statistically significant dishabituation response (investigation greater in trial 4 than trial 3) within each handling method (raw data shown in **Supplementary Fig. 1**). Overall, tail-handled mice demonstrated a non-significant dishabituation response (z = 1.87, P = 0.061), but responses were strain specific: when handled by tail, B6 mice showed a significant response (P = 0.016) but BALB/c mice did not (P = 0.95). Tail-cup-handled mice showed a significant dishabituation response (z = 2.55, P = 0.011) that was consistent across both strains and sexes. Those handled using a tunnel showed a very highly significant dishabituation response (z = 3.18, P = 0.001) that was also consistent across strains and sexes.

We then used the mean and standard deviation of the dishabituation response shown by each handling group in *a posteriori* power calculations to predict the number of independent samples that would be needed to achieve a significant response in future studies, using a significance threshold of p = 0.05 and 80% power to detect a difference (**Table 3**). This predicted that, across all mice, a sample size of n=6 should be sufficient with tunnel handling, while much bigger sample sizes of n=33 would be required for tail-cup handling and n=50 for mice handled by tail. As the effect of handling method depended on strain to some extent, we also predicted required sample sizes for the two strains separately. BALB/c mice were particularly sensitive to handling method for achieving a successful dishabituation response: while a very small sample size of n=6 should achieve 80% power if a handling tunnel is used, this increased to n=35 for tail-cup handling, while BALB/c showed a negative response when handled by tail. Only five B6 mice should be needed if handled by tunnel, while twice this many mice should be needed to achieve 80% power for tail-cup or tail handling.

**Table 3.** Predicted sample size required to achieve 80% power with a = 0.05 based on mean and standard deviation of dishabituation responses.

|  | **Tail** | **Tail-cup** | **Tunnel** |
|---|---|---|---|
| B6 | 9 | 10 | 5 |
| BALB/c | - | 35 | 6 |
| **All mice** | 50 | 33 | 6 |

## Discussion

Our findings indicate that tail-cup handling had some benefits relative to tail handling, with most anxiety measures in the elevated plus maze reduced to a similar level as tunnel handling. Tail-cup experience also reduced avoidance of the handler compared to traditional tail handling, but the willingness of tail-cup-handled mice to interact was still much less than among those handled by tunnel. Tunnel-handled mice also responded much more reliably in the habituation-dishabituation test, such that substantially fewer subjects should be required to show a significant dishabituation response to a novel stimulus, while there was no difference between tail- and tail-cup-handled mice. These differences between methods were consistent across strains and sexes, suggesting that picking up mice using the tail-cup method is better than traditional tail handling, but using a tunnel is still more beneficial both for animal welfare and reliability in behavioural testing.

Two different types of test were used to assess willingness to interact with the handler. The first assessed voluntary interaction with the handling device itself, so measured response to the home cage tunnel held by a handler for tunnel-handled mice or to the gloved hand for tail and tail-cup methods. Consistent with many previous studies ^3,6,7,9,10,14,17,22^, tunnel-handled mice spent substantially more time interacting with their device than tail-handled mice from the first handling session, rising from 6-fold to over 16-fold more time by the ninth handling session. Mice are clearly attracted to spend time interacting with a handling tunnel, despite their experience of being briefly confined and lifted in the tunnel.

This may, in part, be due to a feeling of safety when inside the tunnel. However, mice interact just as readily with an open ladder ^7^ or upturned shelter ^26^ used to pick mice up, so familiarity with a passive object is more likely to be important for stimulating voluntary approach and investigation, even if the device is held in the hand. By contrast, tail-handled mice are initially very reluctant to approach a gloved hand alone, and particularly to touch or climb on the hand, a reluctance that shows little improvement when mice are picked up repeatedly. Mice picked up by the tail-cup method became more willing to interact after repeated experience, suggesting that familiarity with this method helped to overcome an initial aversion. A similar familiarisation process is evident among mice cupped on the open hand without tail restraint, where mice slowly increase their willingness to interact with the hand with increasing experience ^3,9^. After substantial prolonged experience to familiarise animals (60s handling per day over 9 days), unrestrained cup-handled mice show extensive voluntary interaction with the hand, including initiation of paw contact and climbing onto the hand ^3^. We do not know whether tail-cup-handled mice would respond as positively with such extensive handling experience, but this amount of handling is beyond the normal experience of most laboratory mice. With brief daily handling of about 2s, similar to the current study, Gouveia & Hurst ^9^ found mice cupped on the hand without tail restraint showed more voluntary interaction by day 9 than seen here for tail-cup handling, although handler interaction among tunnel and tail groups was very similar to the current study. It is difficult to compare directly between separate studies, but this suggests that willingness to accept contact with the hand by mice cupped on the hand might be slower if combined with tail restraint. However, tail restraint has the advantage that animals cannot jump from the hand if they are not yet habituated to handling. Importantly, though, Gouveia & Hurst ^9^ found that frequent (daily) handling was needed for unrestrained hand-cupped mice to overcome their aversion to the hand and to not run away when approached; neither of these was achieved if animals only experienced this cupped handling at cage cleaning over several weeks. Further research would be needed to establish whether a low frequency of tail-cup handling can be effective in reducing avoidance of the hand or whether additional handling effort is needed to familiarise mice with this first.

To examine a more generalised response to contact with handlers, we also assessed all three handling groups for voluntary interaction with a gloved hand alone. Although this test was only novel for tunnel-handled mice, they still spent much longer interacting with the gloved hand than those handled by tail-cup or tail alone, consistent with previous studies comparing tunnel with tail handling ^3,13,36^. Tail-cup-handled mice again showed an intermediate response that was consistently stronger than tail-handled mice but significantly weaker than those handled by tunnel. Another study by Novak and colleagues ^11^ measured voluntary interaction both with the hand and with a handling tunnel among tail- and tunnel-handled mice. While tunnel-handled BALB/c mice generally interacted more with a hand than those handled by tail, they found no difference among B6 mice and durations were highly variable. However, that study differed from others in that tests were carried out immediately after each other on the same mice, in a different order for each handling method. This prevents direct comparison of equivalent tests between the handling methods and may explain why responses were less clear-cut compared to other studies.

Once familiar with their handling method, we assessed anxiety in a classic EPM test. In agreement with multiple previous studies ^3,6,7,9,10,13,17,22^, mice picked up in a tunnel spent more time on the open arms of the maze and entered more frequently compared to tail-handled mice, both of which are established measures reflecting reduced anxiety ^34,35^. Time spent on the open arms and frequency of entries by tail-cup-handled mice were very similar to tunnel-handled mice and significantly greater than for tail handling. Overall, this suggests that tail-cup and tunnel handling had similar effects on reducing anxiety in this test compared to tail handling. Other studies have shown that cupping on the open hand without restraint ^3,8,9^ or picking up mice on a cage ladder ^7^ similarly reduce anxiety compared to tail handling. While handling method affected both sexes and strains similarly for these two measures, the frequency of protected stretched postures (another EPM anxiety measure) differed only in males. Sex differences in this response to handling method appear to be inconsistent between studies (even in the same laboratory). Comparing tunnel- and tail-handling in B6 mice, Gouveia & Hurst ^9^ found a significantly higher frequency of protected stretch attend only among tail-handled females. However, in the original study by Hurst & West ^3^, tail handling consistently elevated this measure in both sexes of three strains. Overall, taken together with evidence from other EPM measures, picking mice up by the tail appears to promote anxiety consistently in both sexes compared to tunnel, hand-cupping without restraint, or tail-cup methods for handling mice.

Despite lower anxiety in an EPM test using tail-cup compared to tail handling, tail-cup did not improve performance over traditional tail handling in a simple scent discrimination test that was designed to be very easy for mice to perform. Normally, mice of both sexes are strongly motivated to investigate urine scents from other mice, while mice with normal olfactory functioning should easily discriminate between urine from different strains that have very distinct scents ^37^. Nonetheless, our laboratory has shown previously that transferring mice (BALB/c females) to and from a test arena by tunnel or by tail has a major impact on responsiveness, with robust discrimination when handled by tunnel, but failure to discriminate or even investigate the test stimuli when picked up by tail ^21^. Our current study has confirmed robust scent investigation and discrimination among tunnel-handled animals that generalised across sexes and two different strains. Such reliable investigatory responses reduce the number of animals required for testing whilst also enabling more subtle differences between stimuli to be tested. By contrast, animals handled by tail or tail-cup showed substantially weaker responses to unfamiliar male scent, with only a very limited increase in investigation among most mice tested. Gouveia & Hurst ^21^ found that general exploratory behaviour was substantially reduced among tail-handled mice, which strongly predicted poor performance in this dishabituation task. By contrast, mice picked up and transferred by cupping without tail restraint mostly showed good exploratory behaviour and a reasonably strong dishabituation response, albeit with more variation than among tunnel-handled mice. However, tail-cup mice in the current study failed to respond more strongly than tail-handled mice, suggesting that capture and restraint by the tail interfered with performance in this test. The number of replicates that would be needed to achieve a significant response with 80% power was predicted to be very high if mice were handled by either tail or tail-cup due to the combination of small effect size and high variation in these groups; this contrasts markedly with very small numbers required using tunnel handling, where 96% of subjects showed the expected response. This difference was particularly stark in tests using BALB/c mice, which are more responsive to stress and lower risk-taking than B6 ^30–32^. Notably, BALB/c mice responded very reliably if handled using a tunnel. Even the B6 strain required twice as many replicates if handled by tail or tail-cup. This clearly points to the need to use tunnel-handling or an equivalent refined method that does not inhibit responses when conducting this type of behavioural test. More generally, this test reveals that tail and tail-cup handling both disrupt normal exploratory behaviour and motivation to investigate important social information, consistent with negative impacts on animal welfare.

The much poorer performance of tail-cup compared to tunnel-handled mice in the habituation-dishabituation task contrasts with their similar performance in the EPM test. This may be because the two tests assess different aspects of anxiety and exploratory behaviour that are differentially affected by these two handling methods. Open field tests, which are typically conducted in arenas equivalent to that used in the dishabituation test, assess different aspects of anxiety behaviour compared to EPM measures, and also often show contradictory results to EPM tests ^33,38^. However, it is also the case that the habituation-dishabituation test involved several handling events between multiple trials, thus may be more sensitive for revealing impacts caused by negative handling experience. For tail and tail-cup methods, mice had to be captured by their tails from behind in a large open arena, an experience that is imposed on the animals without active cooperation in the procedure. Although the tail-cup method avoids any suspension by the tail, this made little if any difference to their poor habituation-dishabituation response. Nonetheless, over time they developed greater willingness to interact voluntarily with the hand in their home cage, probably because being picked up on the hand allowed animals to become familiar with hand contact ^39^. This was still not as strong as the willingness of all tunnel-handled mice to interact with the hand despite having little prior direct hand contact. The low anxiety, willingness to explore and investigate novel stimuli shown across all three tests by tunnel-handled mice is consistent with previous findings, as discussed above, and adds to strong evidence that this method of picking mice up is very effective for minimising fear and anxiety associated with handling. However, it is important that animals are captured quickly and efficiently, as recommended in training resources ^40^, with tunnel handling taking little or no longer than tail handling once handlers are trained in good technique. Using a different approach to tunnel handling, two studies ^15,16^ failed to see the improvements expected from many other studies. In both cases, the method used for tunnel handling took substantially longer, because handlers waited for animals to enter tunnels voluntarily or did not succeed in getting animals straight into tunnels. As a result, BALB/c mice took more than 5 minutes to enter tunnels voluntarily in one study ^15^, while the time taken to pick up B6 and CD-1 mice in a tunnel was extended by at least fourfold on average compared to tail handling in the other study ^16^. Animals that are not encouraged to enter tunnels immediately, with good technique, experience much greater exposure to disturbance, which appears to remove most of the benefits of tunnel handling and training materials recommend that this should be avoided ^40^.

In conclusion, this study supports the established evidence that use of a tunnel to pick up and transfer mice provides a significant refinement for laboratory mouse welfare compared to traditional tail handling; it can also improve the reliability of behavioural testing, consequently reducing the number of animals required, and improving the robustness of conclusions drawn. Tail-cup handling provides some improvement to welfare over traditional tail handling, helping to overcome aversion to contact with the handler once animals are familiar with the method and reducing some aspects of anxiety associated with being picked up by the tail. However, tail-cup handling failed to improve reliability in our behavioural test and, for most measures in our study, was not as effective a refinement as tunnel handling.

## Methods

The study is reported in line with the ARRIVE guidelines 2.0 ^41^. The essential 10 checklist is available in supplementary materials (**Supplementary Table 5**).

### Ethical Statement

This study was conducted at the University of Liverpool. All tests conducted in this experiment were non-invasive behavioural tests that are not considered to require regulation under the UK Animals (Scientific Procedures) Act 1986. The work was conducted in accordance with the UK Home Office code of practice for the housing and care of animals bred, supplied or used for scientific purposes ^42^. The work was approved by the University of Liverpool AWERB, though no specific licences were required for this study.

### Animals, Housing and Husbandry

The study was conducted using a total of 40 C57BL/6JOlaHsd (B6) and 40 BALB/cOlaHsd (BALB/c) mice, with 20 males and 20 females of each strain, sourced from Envigo, UK. These strains are commonly used as model animals in research making our findings widely generalisable. Upon arrival, mice were aged 3-4 weeks. All mice were handled using a tunnel to transfer them to their home cage before a 7-day undisturbed acclimatisation period, except for daily checks.

Mice were same-sex pair housed in open top M3 cages (48 x 15 x 13cm, North Kent Plastics Rochester, UK) containing Corn Cob Absorb 10/14 substrate (IPS Product Supplies Ltd, London, UK), paper wool nesting material (IPS Product Supplies Ltd), an aspen chew block (1023008, LBS), a clear acrylic tunnel (150mm long□x□50mm wide, Clear plastic supplies, UK), a cardboard tube, and a cardboard nest box. Mice had *ad libitum* access to food (LabDiet 5LF2, EURodent Diet 14%, IPS Product Supplies Ltd) and water at all times in their home cages. Cages were cleaned every two weeks, and mice were handled using their assigned handling method during transfer to clean cages. Mice were maintained on a reverse 12:12 light/dark cycle, with white lights on between 20:00 and 08:00. All handling and testing were conducted under red light during the first four hours of the dark phase when mice are most active. Animals were kept at a constant temperature (20-24°C) and humidity (45-65%). The animal’s tails were marked for identification every two weeks using an animal marker, until PIT tags were implanted for long-term identification at either 19 or 26 days after the first handling session.

Each cage-pair was randomly assigned to one of three handling groups by random number generator (Excel’s RAND function): tunnel (n=24 mice/12 cages), tail (n=24 mice/12 cages), or tail-cup (n=24 mice/12 cages), balanced by sex and strain. These sample sizes were calculated using G*Power (version 3.1.9.6) ^43^ based on effect sizes observed between tail- and tunnel-handled mice in previous experiments within our lab ^3,21^. An additional 8 mice (two per sex and strain combination) were obtained at the same time to allow for any withdrawal from the study. These animals were randomly allocated to handling groups, adding one more pair of B6 males to tunnel handling and the other three pairs to tail handling. Mice were handled exclusively by their assigned handling method during husbandry and testing procedures. Cage locations for handling treatments were balanced on the home cage rack. The first experimental handling session and testing began when mice were 8-11 weeks old, having experienced handling twice during cage cleans prior to the formal start of testing. Where evidence of fighting was observed, the pair were split (n=3 pairs). During the study, three mice were humanely killed due to health concerns. Resulting sample sizes are shown in supplementary materials (**Supplementary Table 6**).

### Study Design and Timeline

As all mice could not be tested on the same day, the elevated plus maze (EPM) and habituation-dishabituation tests were staggered over multiple days, with each mouse tested only once for each test (**Figure 4**). Behaviours were scored from video recordings, blind to handling methods where possible. This included the EPM, Voluntary Interaction Tests (VITs) with gloved hand, and habituation-dishabituation trials. VITs with the handling device could not be blinded for tunnel handling as interaction with the handling device was the main outcome measure. However, comparison between tail and tail-cup groups was scored blind, as interaction for both was with a gloved hand. Testing orders were randomised and counterbalanced for handling method, sex and strain throughout.

**Figure 4.**
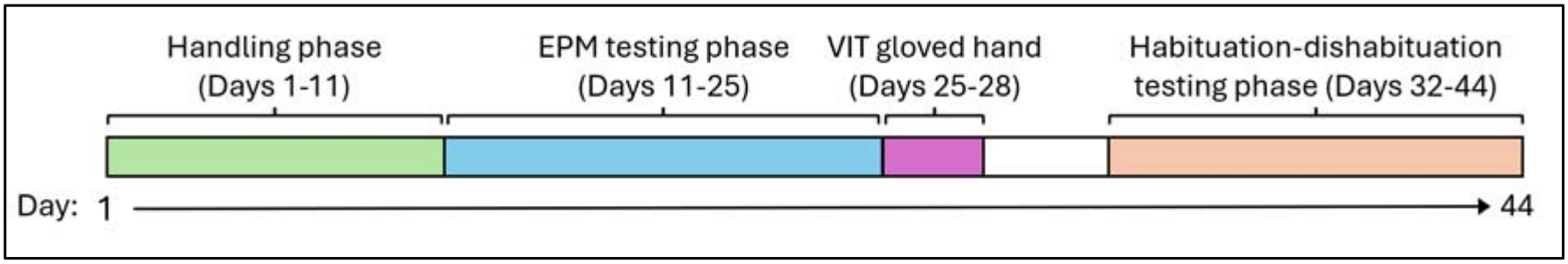
Study timeline showing when each of the tests was conducted, relative to the first handling session (day 1). Tests were conducted on multiple days, but each mouse was used only once in each test. Voluntary interaction tests were conducted after the first and ninth (final) handling session during the handling phase.

### Daily Handling Sessions and Voluntary Interaction Tests (VIT)

Mice experienced nine daily handling sessions, when they were picked up briefly (approximately 2s) by their assigned handling method: tail, tunnel or tail-cup. Prior to each handling session, the contents of the home cage were removed, leaving only bedding substrate. Tail-handled animals were picked up by the base of their tail, using the thumb and forefinger, and lifted briefly ^4,5^. For tunnel-handled animals, the clear acrylic tunnel in their home cage was held in front of the animal, which they either immediately entered voluntarily or were gently encouraged straight into with the other hand and the tunnel lifted with the mouse inside ^3^. The ends of the tunnel did not need to be covered. For mice handled by tail-cup, the base of the tail was held between the thumb and forefinger before scooping the hand underneath the animal so that it sat on the open palm before lifting.

Voluntary interaction tests to measure the willingness of animals to interact with the handling device were conducted following the first and last handling sessions. After handling, the experimenter stood motionless in front of the open cage for 60 s. The handling device (gloved hand for tail- and tail-cup-handled mice, and gloved hand holding home cage tunnel for tunnel-handled mice) was held in the front half of the cage nearest the experimenter for 60 s. The time that mice spent voluntarily interacting with the handling device was scored from overhead video recordings using BORIS software (version 7.10.7) ^44^. Interaction was considered as any physical contact or nose within 0.5cm of the handling device. As mice within the same cage were tested together, total interaction time was averaged across mice in the same cage, and data were analysed at the cage level.

A voluntary interaction test to compare the willingness of animals to interact with a gloved hand alone followed the same method, but in this case a gloved hand alone was introduced into the home cage for all treatment groups for 60 s. This test was conducted the day following the EPM test.

### Elevated Plus Maze

Anxiety levels were assessed in an EPM test. Two smooth white plastic mazes had two open and two closed arms (each 30 x 5cm), raised 58cm from the ground. The closed arms had walls 15cm in height. Mice were delivered to the centre of the EPM by their assigned handling method, facing towards a closed arm. Mice in each cage pair were tested simultaneously on separate mazes with a visual barrier placed between the two. Trials lasted five minutes and were recorded by camera from directly above. Time spent on each arm, number of entries, and number of protected stretch attend postures were recorded using BORIS software. Mazes were cleaned with 70% ethanol and thoroughly dried between trials.

### Habituation-Dishabituation

Methods followed those used by Gouveia & Hurst ^21^. Mice from each pair were tested simultaneously in two identical clean arenas (L70cm x W60cm x H55cm, white laminated MDF). Each mouse experienced four 5-minute trials, with five-minute inter-trial intervals when the mouse was removed to an empty holding cage (M3) and the test arena and plastic tile were cleaned with 70% ethanol and thoroughly dried. Mice were transferred to and from the arena using their assigned handling method and placed at the edge of the arena at the start of a trial.

In each trial, a urine stimulus (10μl male mouse urine) was presented as a single streak on a clean glass microscope slide secured to a clear acrylic tile (14.5 x 14.5cm), placed in the centre of the arena. For the first three trials (habituation phase), the stimulus came from a different strain from the subject so that it was initially unfamiliar. On the fourth trial (dishabituation), urine came from a third strain (different from the subject and the habituation stimulus) so that it was again unfamiliar and chosen such that mice should easily be able to tell the difference ^37^. To achieve this, urine samples were collected from male donors of three laboratory strains: ICR (n=4), BALB/c (n=11), and B6 (n=5). Donors were adult stock animals, aged at least three months and housed singly so that all were of equivalent social status. Urine was collected in standard MB1 cages (45 x 28 x 13cm) fitted with metal grid floors for up to 30 minutes during the first two hours of the active phase of the light cycle. Urine from each strain was pooled and frozen at −20°C for up to four weeks before use, with individual aliquots defrosted for each set of trials to avoid repeated freeze-thawing of a larger pool. The strain used as the habituation or dishabituation stimulus was swapped between mice in the same cage pair; for example, for B6 mice, one received BALB/c urine for trials 1-3 and ICR for trial 4, while its cagemate received these the other way round.

Trials were video recorded from directly above, and the total time spent investigating the scent stimulus during each trial (nose within 0.5cm of the urine-streaked tile) was scored from recordings using a Karrimor stopwatch.

### Data Analysis

Analyses were conducted in RStudio version 4.3.2 ^45^. General linear models used the lm() function from the “stats” package, and general linear mixed effects models were fitted using the “lme4” and “lmerTest” packages ^46,47^. Following model fitting, normality of model residuals was visually inspected using histograms and Q-Q plots, and tested using a Shapiro-Wilk test. Where necessary, data were transformed using either log or square root transformations to achieve normal distribution of residuals.

Model types, data transformations, terms included in each model and sample sizes are listed in **Supplementary Table 6**. As mice were pair housed and handling treatment applied at cage level, cage was included as a random factor in models to avoid potential pseudoreplication ^48,49^, except for voluntary interaction tests where time interacting was averaged over mice in the same cage and analysed with cage as the experimental unit.

However, where the random factor variance was equal or near zero, the random factor was removed to allow non-singular model fit ^50^. Significance criterion for p-values was considered < 0.05. For each model, interaction terms were initially included between all main factors. Non-significant interaction terms were then removed from final models ^51^. Interaction terms that neared significance (0.05 < p < 0.1) and so were suggestive but inconclusive of a possible effect were included ^52^. P-values from models were obtained from ANOVA tables with Satterthwaite’s method. Where a main effect of handling method was found, the emmeans() function from the “emmeans” package ^53^ with Tukey adjustment for multiple testing was used to assess post hoc pairwise comparisons between the three methods.

For dishabituation responses, parametric model residuals showed high levels of kurtosis, so non-parametric analyses assessed the mean dishabituation response for each cage. The difference in investigation between trial 4 minus trial 3 was compared between handling methods with a Kruskal-Wallis test, while pairwise comparisons between each pair of handling methods were then assessed using Dunn tests adjusted with a Bonferroni correction for multiple comparisons. Wilcoxon matched-pair signed ranks tests compared trial 4 to trial 3 to assess the significance of a dishabituation response within each handling method. These non-parametric tests provided the same conclusions as the original linear mixed-effect models, but the latter are not reported here.

Using the results of the habituation-dishabituation test, we conducted *a posteriori* power calculations to predict the number of independent samples required for each individual handling method to see a significant dishabituation response with 80% power and p < 0.05. These were conducted using G*Power (version 3.1.9.6) ^43^ based on a matched-pairs t-test. Effect size (Cohen’s d) was calculated as the mean dishabituation response observed / standard deviation of dishabituation response for each handling group.

### Missing or Excluded Data

Videos for two trials were lost for the VIT following the ninth handling session due to recording error and a technical issue. Two trials were lost from the EPM analysis, as mice jumped off the maze.

Where data points were potentially considered outliers based on graphical presentation of residuals in Q-Q plots, we repeated analyses with outliers excluded to assess the impact on our conclusions. In all cases, this had no effect on the conclusions drawn and the original analyses including all data points are reported.

Faecal samples were collected from subjects 4 hours following their EPM test (peak corticosterone response following increased adrenocortical activity ^54^) with the aim of investigating the effects of handling method on hypothalamic-pituitary-adrenocortical stress reactivity. However, a valid assay could not be established due to problems with consistent extraction of corticosterone metabolites, so these data are not reported.

## Acknowledgements

We would like to thank Amanda Davidson, Olivia Whittle and Rick Humphries for technical support during the study, along with the Mammalian Behaviour and Evolution group for constructive feedback at all stages of the project. We thank Jonathan Nolan for help in construction of the elevated plus mazes. The work was funded by the NLD (Newcastle-Liverpool-Durham) BBSRC Doctoral Training Partnership, held by R.N.

## Author contributions

J.L.H., M.L. & C.R. gained funding for the studentship; J.L.H., M.L. & C.R. provided supervision; R.N. & J.L.H. designed the study and protocols; R.N. carried out the experiments; R.N. & J.L.H. analysed the data and drafted the manuscript; M.L. & C.R. commented on the manuscript.

## Competing Interests Statement

R.N. was employed by National Centre for the Replacement, Refinement and Reduction of Animals in Research (NC3Rs) during preparation of the manuscript. The authors declare no other competing interests.

## Supplementary Materials

See PDF file.

