## Supplementary materials for "The impact of tail-cup handling as an alternative method for picking up laboratory mice"

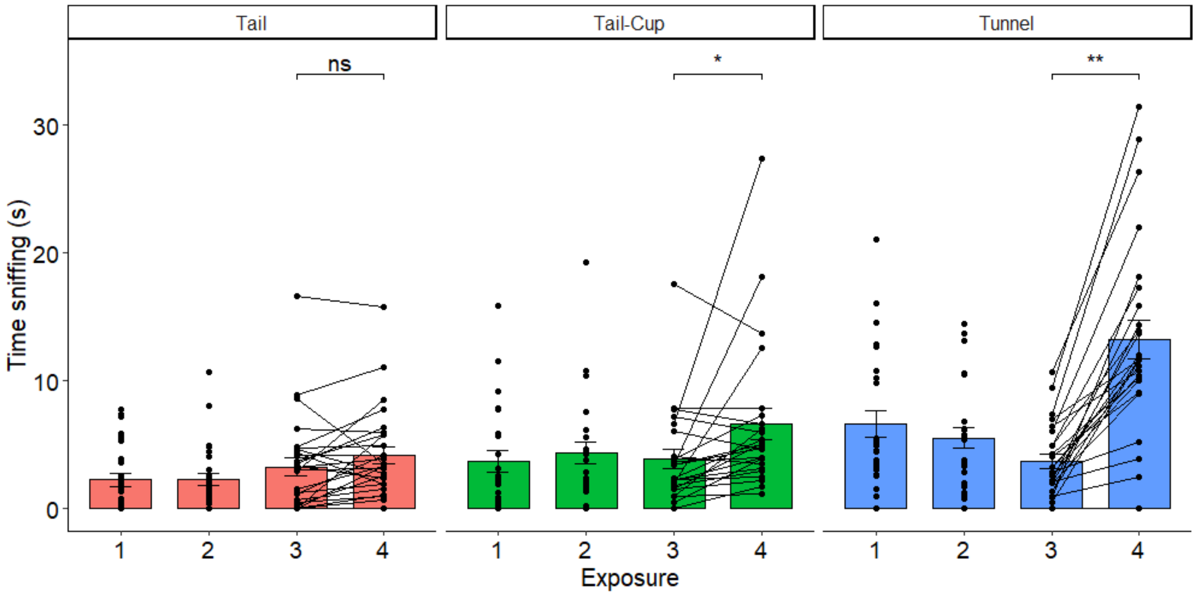


**Supplementary Figure 1**. Stimulus investigation in a habituation-dishabituation task for different handling methods, pooled across sex and strains (mean ± SEM). A decrease between trial 1 and 3 represents the habituation phase as mice become familiar with the same urine presented in all three trials. However, low investigation in trial 1 is typical when mice are unfamiliar with the test arena (Gouveia & Hurst [1]). An increase between trials 3 and 4 represents the dishabituation response, where mice increase investigation of an unfamiliar urine stimulus in trial 4. Points for the same individual are joined by lines between trials 3 and 4, to visualise the response. P values from Wilcoxon matched-pair signed rank tests: ns P > 0.05, *P < 0.05, **P < 0.01.


1. Gouveia, K. & Hurst, J. L. Optimising reliability of mouse performance in behavioural testing: the major role of non-aversive handling. *Sci. Rep.* **7**, 44999 (2017).

**Supplementary Table 1**. Full models including interactions for effects of handling method, sex, and strain on willingness to interact voluntarily with the handler (GLM ANOVAs).

| Voluntary Interaction | Handling Session 1 | | Handling Session 9 | | Gloved Hand | |
|---|---|---|---|---|---|---|
| Handling | $F_{2,29} = 37.69$ | **P < 0.0001** | $F_{2,27} = 90.98$ | **P < 0.0001** | $F_{2,30} = 39.74$ | **P < 0.0001** |
| Sex | $F_{1,29} = 0.15$ | P = 0.70 | $F_{1,27} = 0.57$ | P = 0.46 | $F_{1,30} = 0.18$ | P = 0.68 |
| Strain | $F_{1,29} = 1.05$ | P = 0.31 | $F_{1,27} = 3.70$ | P = 0.065 | $F_{1,30} = 1.77$ | P = 0.19 |
| Handling x Sex | $F_{2,29} = 2.71$ | P = 0.08 | $F_{2,27} = 0.60$ | P = 0.55 | $F_{2,30} = 2.05$ | P = 0.15 |
| Handling x Strain | $F_{2,29} = 0.17$ | P = 0.84 | $F_{2,27} = 0.98$ | P = 0.39 | $F_{2,30} = 0.30$ | P = 0.74 |
| Sex x Strain | $F_{1,29} = 0.55$ | P = 0.46 | $F_{1,27} = 0.82$ | P = 0.37 | $F_{1,30} = 0.95$ | P = 0.34 |
| Handling x Sex x Strain | $F_{2,29} = 0.29$ | P = 0.75 | $F_{2,27} = 0.88$ | P = 0.42 | $F_{2,30} = 0.82$ | P = 0.45 |

**Supplementary Table 2**. Full models including interactions for effects of handling method, sex, and strain on open arm duration and entries during the elevated plus maze test (GLMM ANOVAs).

| Elevated Plus Maze | Duration open arms | | No. entries open arms | |
|---|---|---|---|---|
| *Full Model* | | | | |
| Handling | $F_{2,29.4} = 7.66$ | **P = 0.002** | $F_{2,27.4} = 6.01$ | **P = 0.007** |
| Sex | $F_{1,29.4} = 0.97$ | P = 0.33 | $F_{1,27.4} = 0.03$ | P = 0.86 |
| Strain | $F_{1,29.4} = 22.30$ | **P < 0.0001** | $F_{1,27.4} = 4.61$ | **P = 0.041** |
| Handling x Sex | $F_{2,29.4} = 1.30$ | P = 0.29 | $F_{2,27.4} = 1.91$ | P = 0.17 |
| Handling x Strain | $F_{2,29.4} = 0.52$ | P = 0.60 | $F_{2,27.4} = 1.73$ | P = 0.20 |
| Sex x Strain | $F_{1,29.4} = 6.04$ | **P = 0.02** | $F_{1,27.4} = 6.30$ | **P = 0.018** |
| Handling x Sex x Strain | $F_{2,29.4} = 1.11$ | P = 0.34 | $F_{2,27.4} = 2.04$ | P = 0.15 |

**Supplementary Table 3**. Full models including interactions for effects of handling method, sex, and strain on protected stretch attend postures during the elevated plus maze test (GLMM ANOVAs).

| Elevated Plus Maze | No. PSA | | No. PSA (Females) | | No. PSA (Males) | |
|---|---|---|---|---|---|---|
| *Full Model* | | | | | | |
| Handling | $F_{2,26.9} = 3.97$ | **P = 0.031** | $F_{2,14.4} = 0.89$ | P = 0.43 | $F_{2,9.8} = 9.31$ | **P = 0.005** |
| Sex | $F_{1,26.9} = 2.76$ | P = 0.11 | | | | |
| Strain | $F_{1,26.9} = 4.12$ | **P = 0.052** | $F_{1,14.4} = 2.99$ | P = 0.11 | $F_{1,10.0} = 1.14$ | P = 0.31 |
| Handling x Sex | $F_{2,26.9} = 3.33$ | **P = 0.051** | | | | |
| Handling x Strain | $F_{2,26.9} = 2.08$ | P = 0.14 | $F_{2,14.4} = 0.68$ | P = 0.52 | $F_{2,9.8} = 2.14$ | P = 0.17 |
| Sex x Strain | $F_{1,26.9} = 0.60$ | P = 0.44 | | | | |
| Handling x Sex x Strain | $F_{2,26.9} = 0.11$ | P = 0.90 | | | | |
| *Final Model (non-significant interactions removed)* | | | | | | |
| Handling | $F_{2,32.9} = 3.72$ | **P = 0.035** | $F_{2,16.4} = 0.95$ | P = 0.41 | $F_{2,14.4} = 8.11$ | **P = 0.004** |
| Sex | $F_{1,33.0} = 3.44$ | **P = 0.072** | | | | |
| Strain | $F_{1,32.9} = 3.83$ | **P = 0.059** | $F_{1,16.5} = 2.60$ | P = 0.13 | $F_{2,14.3} = 1.18$ | P = 0.30 |
| Handling x Sex | $F_{2,33.0} = 3.41$ | **P = 0.045** | | | | |
| Post hoc comparisons: | | | | | | |
| Tail vs Tail-Cup | | | | | | P = 0.53 |
| Tail vs Tunnel | | | | | | **P = 0.003** |
| Tunnel vs Tail-Cup | | | | | | **P = 0.039** |

**Supplementary Table 4**. Effects of handling method, sex, and strain on control measures (closed arm entries and duration) during the elevated plus maze test (ANOVAs).

| **Elevated Plus Maze** | **Duration closed arms** | | **No. entries closed arms** | |
|---|---|---|---|---|
| *Full Model* | | | | |
| Handling | $F_{2,64} = 0.27$ | $P = 0.76$ | $F_{2,64} = 0.43$ | $P = 0.65$ |
| Sex | $F_{1,64} = 0.54$ | $P = 0.46$ | $F_{1,64} = 0.33$ | $P = 0.57$ |
| Strain | $F_{1,64} = 21.98$ | **$P < 0.0001$** | $F_{1,64} = 32.54$ | **$P < 0.0001$** |
| Handling x Sex | $F_{2,64} = 0.91$ | $P = 0.41$ | $F_{2,64} = 0.41$ | $P = 0.66$ |
| Handling x Strain | $F_{2,64} = 0.88$ | $P = 0.42$ | $F_{2,64} = 1.29$ | $P = 0.28$ |
| Sex x Strain | $F_{1,64} = 1.45$ | $P = 0.23$ | $F_{1,64} = 1.92$ | $P = 0.17$ |
| Handling x Sex x Strain | $F_{2,64} = 0.78$ | $P = 0.46$ | $F_{2,64} = 0.35$ | $P = 0.71$ |
| *Final Model (non-significant interactions removed)* | | | | |
| Handling | $F_{2,71} = 0.27$ | $P = 0.76$ | $F_{2,71} = 0.44$ | $P = 0.65$ |
| Sex | $F_{1,71} = 0.54$ | $P = 0.46$ | $F_{1,71} = 0.33$ | $P = 0.57$ |
| Strain | $F_{1,71} = 22.11$ | **$P < 0.0001$** | $F_{1,71} = 33.0$ | **$P < 0.0001$** |

**Supplementary Table 5**. Author checklist for ARRIVE 2.0 essential 10.

### The ARRIVE guidelines 2.0: author checklist

#### The ARRIVE Essential 10

These items are the basic minimum to include in a manuscript. Without this information, readers and reviewers cannot assess the reliability of the findings.

| Item | | Recommendation | Section/line number, or reason for not reporting |
|---|---|---|---|
| **Study design** | 1 | For each experiment, provide brief details of study design including: | |
| | | a. The groups being compared, including control groups. If no control group has been used, the rationale should be stated. | Introduction & Methods: Animals, Housing and Husbandry section |
| | | b. The experimental unit (e.g. a single animal, litter, or cage of animals). | Methods: data analysis section & supplementary table 6 |
| **Sample size** | 2 | a. Specify the exact number of experimental units allocated to each group, and the total number in each experiment. Also indicate the total number of animals used. | Methods: Animals, Housing and Husbandry section |
| | | b. Explain how the sample size was decided. Provide details of any *a priori* sample size calculation, if done. | Methods: Animals, Housing and Husbandry section |
| **Inclusion and exclusion criteria** | 3 | a. Describe any criteria used for including and excluding animals (or experimental units) during the experiment, and data points during the analysis. Specify if these criteria were established *a priori*. If no criteria were set, state this explicitly. | Methods: Excluded animals section. These were not determined a priori |
| | | b. For each experimental group, report any animals, experimental units or data points not included in the analysis and explain why. If there were no exclusions, state so. | Methods: Excluded animals section |
| | | c. For each analysis, report the exact value of *n* in each experimental group. | Supplementary table 6 |
| **Randomisation** | 4 | a. State whether randomisation was used to allocate experimental units to control and treatment groups. If done, provide the method used to generate the randomisation sequence. | Methods: Animals, Housing and Husbandry section |
| | | b. Describe the strategy used to minimise potential confounders such as the order of treatments and measurements, or animal/cage location. If confounders were not controlled, state this explicitly. | Methods: Animals, Housing and Husbandry section |
| **Blinding** | 5 | Describe who was aware of the group allocation at the different stages of the experiment (during the allocation, the conduct of the experiment, the outcome assessment, and the data analysis). | Methods: Study Design and Timeline |
| **Outcome measures** | 6 | a. Clearly define all outcome measures assessed (e.g. cell death, molecular markers, or behavioural changes). | Supplementary table 6 |
| | | b. For hypothesis-testing studies, specify the primary outcome measure, i.e. the outcome measure that was used to determine the sample size. | Methods: Animals, Housing and Husbandry section |
| **Statistical methods** | 7 | a. Provide details of the statistical methods used for each analysis, including software used. | Methods: Data Analysis section & supplementary table 6 |
| | | b. Describe any methods used to assess whether the data met the assumptions of the statistical approach, and what was done if the assumptions were not met. | Methods: Data Analysis section |
| **Experimental animals** | 8 | a. Provide species-appropriate details of the animals used, including species, strain and substrain, sex, age or developmental stage, and, if relevant, weight. | Methods: Animals, Housing and Husbandry section |
| | | b. Provide further relevant information on the provenance of animals, health/immune status, genetic modification status, genotype, and any previous procedures. | Methods: Animals, Housing and Husbandry section |
| **Experimental procedures** | 9 | For each experimental group, including controls, describe the procedures in enough detail to allow others to replicate them, including: | |
| | | a. What was done, how it was done and what was used. | Methods |
| | | b. When and how often. | Methods |
| | | c. Where (including detail of any acclimatisation periods). | Methods |
| | | d. Why (provide rationale for procedures). | n/a |
| **Results** | 10 | For each experiment conducted, including independent replications, report: | |
| | | a. Summary/descriptive statistics for each experimental group, with a measure of variability where applicable (e.g. mean and SD, or median and range). | Figures 1, 2 and 3 |
| | | b. If applicable, the effect size with a confidence interval. | n/a |

**Supplementary Table 6**. The models/tests and associated information for each of the tests conducted in this study.

| Test | Outcome Variable | Transformation | Model/Statistical Test | Fixed Effects | Random Effects | Test Unit | **Sample Size** (Split evenly between strains and sexes, unless otherwise stated) |
|---|---|---|---|---|---|---|---|
| Voluntary Interaction Test (Handling session 1 and 9) | Proportion of time spent interacting with handling device during 60s test | Square root | General Linear Model (ANOVA) | Handling treatment (3 levels), Strain (2 levels), Sex (2 levels) | - | Cage | **n=15 Tail** (8xBALB/c*, 7xB6(3m/4f)) **n=13 Tunnel** (6xBALB/c, 7xB6(4m/3f)) **n=12 Tail-Cup** |
| | Proportion of time spent interacting with handling device (session 1 vs 9) | Square root | General Linear Mixed Model (ANOVA) | Session (2 levels), Strain (2 levels), Sex (2 levels) | Cage | Cage | **n=15 Tail** (6xBALB/c (4m*/2f†), 7xB6(3m/4f))<br>**n=13 Tunnel** (6xBALB/c, 7xB6(4m/3f))<br>**n=12 Tail-Cup** |
| Elevated Plus Maze | Time on open arms | - | General Linear Mixed Model (ANOVA) | Handling treatment (3 levels), Strain (2 levels), Sex (2 levels) | Cage | Mouse | **n=27 Tail** (14xBALB/c, 13xB6(5m†/8f) **n=25 Tunnel** (12xBALB/c, 13xB6(8m/5f†)) **n=24 Tail-Cup** |
| | Number of entries onto open arms | Log | | | | | |
| | Number of protected stretch attend postures | Square root | | | | | |
| | Control measures – entries to and duration on closed arms | - | General Linear Model (ANOVA) | | -*** | Mouse | |
| Voluntary Interaction Test (Gloved hand) | Proportion of time spent interacting with gloved hand during 60s test | Square root | General Linear Model (ANOVA) | Handling treatment (3 levels), Strain (2 levels), Sex (2 levels) | - | Cage | **n=15 Tail** (8xBALB/c**, 7xB6(3m/4f) **n=13 Tunnel** (6xBALB/c, 7xB6(4m/3f)) **n=12 Tail-Cup*** |
| Habituation-Dishabituation Test | Dishabituation between handling methods | - | Kruskal-Wallis test | Handling treatment (3 levels) | - | Cage | **n=15 Tail** (8xBALB/c**, 7xB6(3m/4f) **n=13 Tunnel** (6xBALB/c, 7xB6(4m/3f)) **n=12 Tail-Cup*** |
| | Dishabituation (Time sniffing) between exposure 3 and 4 | | Wilcoxon matched-pair signed ranks tests | Trial number (2 levels) | - | | |

*One of male cages split (not reflected in number).
† Data lost due to video/test error.
**One of male cages split (not reflected in number). One male and one female cage housed singly due to culling of partner.
*** Cage not included to allow non-singular fit of model, when variance = 0. See Methods section for details.
